# Native catalase expression in *Arabidopsis thaliana* is more than sufficient to limit excess decarboxylation from photorespiratory intermediates

**DOI:** 10.64898/2025.12.06.692765

**Authors:** Luke M. Gregory, Kate F. Scott, Faith Twinamanni, Deserah D. Strand, Han Bao, Andreas P. M. Weber, Berkley J. Walker

## Abstract

The hydrogen peroxide (H_2_O_2_) scavenging enzyme, catalase, plays a critical role in the photorespiratory pathway by maintaining the balance of H_2_O_2_, a reactive oxygen species (ROS), in the peroxisome. H_2_O_2_ acts as both a signaling molecule and a potential source of ROS depending on its accumulation in the peroxisome. Additionally, H_2_O_2_ can also drive non-enzymatic decarboxylation (NED) reactions as well as other decarboxylation reactions, leading to increased CO_2_ release that is linked to a severe growth phenotype. However, the exact cause of this stunted growth phenotype is not fully understood, and it remains unclear whether the capacity of catalase is critical for minimizing these decarboxylating reactions. Here we elucidate the mechanism behind the decrease in plant growth due to the accumulation of H_2_O_2_ from photorespiration using *cat2* knock-out lines of *Arabidopsis thaliana* rescued with transgenic expression lines of *Heliobacter pylori* catalase. These experiments demonstrated that while one of the three heterologous lines expressing *H. pylori* catalase isoform (Hp615) had greater catalase activity than *cat2*-KO and rescued the severe growth and photosynthetic phenotype, its catalase activity was still far below wild type levels. These findings suggest that catalase plays a crucial role in maintaining H_2_O_2_ homeostasis within the peroxisome and minimizing decarboxylation reactions, both of which are linked to plant growth. Moreover, once a threshold capacity is reached, increasing catalase capacity further may offer limited benefits in enhancing net carbon fixation.

**Highlight:** We show that peroxisomal catalase is important for maintaining high rates of net carbon fixation associated with plant growth. Native catalase levels in *Arabidopsis thaliana* are in excess of that which is required to minimize alternative decarboxylation reactions. Therefore, efforts to optimize catalase-mediated degradation of H_2_O_2_ may be of limited benefit.

## Introduction

Plant growth is closely linked to net carbon fixation (*A*), which can be accurately represented using biochemical models based on the reaction kinetics of ribulose 1,5,-bisphosphate carboxylase/oxygenase (rubisco) for either CO_2_ or O_2_ in C_3_ photosynthetic plants (von Caemmerer, 2000; von Caemmerer *et al*., 1981; Walker *et al*., 2016). Carboxylation of ribulose 1,5,-bisphosphate (RuBP) by rubisco (*v*_*c*_) leads to a gain in *A*, while oxygenation (*v*_*o*_) of RuBP initiates photorespiration with the production of 2-phosphoglycolate (2-PG) resulting in a reduction of *A* (Bowes *et al*., 1971). The decline in *A* from photorespiration is due to the release of CO_2_ involved in recycling of 2-PG and the decrease in rubisco efficiency whereby every oxygenation of RuBP is a missed opportunity to carboxylate RuBP (Sharkey, 1988). Due to these inefficiencies, photorespiration has been a critical area of study for improving *A*, and thus plant growth, in C_3_ plants. This realization has led to the development of mutants in key photorespiratory enzymes to explore how perturbations in photorespiration alter plant growth (Somerville, 1984; Somerville *et al*., 1979).

Several knockout mutants of photorespiratory enzymes have been characterized to determine how insufficient conversion of photorespiratory intermediates limit plant growth. The photorespiratory pathway consists of eleven enzymes – rubisco, phosphoglycolate phosphatase, glycolate oxidase, catalase, (glutamate, alanine, serine): glyoxylate aminotransferases, glycine decarboxylase complex, serine hydromethyltransferase, hydroxypyruvate reductase, and glycerate kinase – that work together in a series of linked reactions to detoxify 2-PG and recycle its carbon back to 3-phosphoglycerate. Among the studies on photorespiratory knockout mutants, numerous have identified several pathway intermediates as biologically inert - having little-to-no apparent effect on surrounding biological processes - or biologically active, significantly impacting growth and development due to their accumulation. The first intermediate, 2-PG, the substrate for phosphoglycolate phosphatase, inhibits key C_3_ cycle enzymes including triose phosphate isomerase and sedoheptulose 1,7-bisphosphate (Anderson, 1971; Flügel *et al*., 2017). Accumulation of 2-PG also sequesters inorganic phosphate, reducing its availability for ATP synthesis in the light reactions. This reduction in ATP supply constrains the C_3_ cycle, and limits *A* (Harley *et al*., 1991; Sharkey, 1985; Timm *et al*., 2019). Similarly, glycolate, the substrate for glycolate oxidase may inhibit rubisco, which reduces A (Cui *et al*., 2016; González-Moro *et al*., 1997; Lu *et al*., 2014; Wendler *et al*., 1992; Xu *et al*., 2009). Hydrogen peroxide (H_2_O_2_) is another biologically active intermediate that serves as a signaling molecule, but when it accumulates, is a source of oxidative stress and reduces plant growth thought to be due in part to its participation in decarboxylation reactions (Bao *et al*., 2021; Cousins *et al*., 2008; Dat *et al*., 2003; Grodzinski, 1978; Halliwell *et al*., 1974; Keech *et al*., 2012; Mittler, 2017; Springsteen *et al*., 2018). These explorations of photorespiratory knockout mutants have negatively correlated plant growth to the accumulation of biologically active intermediates which accumulate when photorespiration is perturbed.

Among the biologically active photorespiratory intermediates, H_2_O_2_ stands out due to its role as a reactive oxygen species (ROS) and signaling molecule, with photorespiration serving as a major metabolic source of its production in the light at least outside the chloroplast (Foyer *et al*., 2003). H_2_O_2_ is produced when glycolate and H_2_O is converted to glyoxylate and H_2_O_2_ in the peroxisome by glycolate oxidase. While glyoxylate continues through the pathway and ultimately converted to 3-phosphoglycerate and reenters the Calvin-Benson cycle, H_2_O_2_ remains in the peroxisome. The increase in endogenous H_2_O_2_ in the light can trigger various developmental and physiological processes including programmed cell-death (Cheng *et al*., 2016; Vavilala *et al*., 2015), stomatal aperture regulation and movement (Ge *et al*., 2015; Rodrigues *et al*., 2022; Shi *et al*., 2024), decarboxylation reactions (Bao *et al*., 2021; Cousins *et al*., 2008; Grodzinski, 1978; Halliwell *et al*., 1974; Keech *et al*., 2012), and as a signaling molecule in abiotic stress (Ashraf *et al*., 2015; Hameed *et al*., 2014; Wang *et al*., 2014; Wu *et al*., 2015). While all these physiological processes are important, H_2_O_2_ participation in decarboxylation reactions may directly limit plant growth, making it a key target for metabolic engineering. Regulating H_2_O_2_ levels within the peroxisome is therefore essential for reducing excess CO_2_ release and mitigating the broader physiological impacts of ROS overaccumulation.

Early signs of catalase controlling H_2_O_2_ concentration were in the identification of chemically mutagenized seeds in *Arabidopsis thaliana*, which failed to grow under ambient CO_2_ concentrations, but had normal growth at high CO_2_-enriched air (Somerville et al., 1982). In a similar study, *Hordeum vulgare L*. mutants that failed to grow in air exhibited 90% loss in catalase activity compared to parents in the F2 and F3 generations (Kendall et al., 1983). Later, a population of haploid *Nicotiana tabacum* L. cv Havana Seed plantlets were screened under high O_2_ and low CO_2_ treatment (42% O_2_ and 160 ppm CO_2_) and selected for O_2_-resistance. The O_2_-resistant *N. tabacum* were found to have 40%-50% greater catalase activity than wildtype (WT) lines (Zelitch, 1989, 1992). Additionally, these high catalase lines had a corresponding increase in A, suggesting that H_2_O_2_ plays a role in photorespiratory CO_2_ release. This brings up the question whether catalase activity is in excess to suppress these reactions, or whether these decarboxylation reactions occur normally.

The peroxisome-localized catalase knockout (*cat2-*KO) mutant has emerged as a valuable model for investigating the role of catalase and the balance between H_2_O_2_ production and scavenging in photosynthetic tissue (Bao *et al*., 2021; Kendall *et al*., 1983; Queval *et al*., 2007). The stoichiometric loss of CO_2_ per oxygenation in *Arabidopsis thaliana cat2-*KO lines is greater than WT lines at 25°C, due to the increased frequency of decarboxylation reactions (0.64 and 0.5, respectively) (Bao *et al*., 2021). The lower photorespiratory efficiency in *cat2*-KO, defined here as the carbon recycling efficiency of photorespiration, reduces *A* and plant growth compared to WT lines. The lower photorespiratory efficiency in *cat2*-KO compared to WT appears to be due to the deficiency in the maximal reaction velocity (*V*_*max*_) of catalase, which is unable to keep pace with the photorespiratory demand (*v*_*o*_). Although these decarboxylation reactions occur more frequently in *cat2*-KO lines due to the lack of catalase capacity, it is unclear whether catalase activity is in excess in WT lines to eliminate the decarboxylation reactions.

Replacing native *A. thaliana* catalase with a more efficient isoform has the potential to improve photorespiratory efficiency if the isoform can reduce CO_2_ release per oxygenation below the WT stoichiometry. If lower concentrations of H_2_O_2_ were maintained in the peroxisome, we would expect a decline in NED due to the first-order reaction kinetics of these reactions. To investigate this question, we generated three transgenic independent expression lines of *Heliobacter pylori* catalase transformed in *A. thaliana cat2*-KO to determine their *in vivo* and *in vitro* impact to photosynthetic and photorespiratory capacity. *H. pylori* catalase was selected based on;’. *H. pylori* has a lower Michalis-Menten constant (*K*_*m*_; 43-127 mM) compared to *A. thaliana* (138 mM) indicating a higher affinity for H_2_O_2_, thus increasing H_2_O_2_ detoxification in the peroxisome for a given *V*_*max*_ (Hazell *et al*., 1991; Switala *et al*., 2002).

Our transgenic rescues of *cat2-*KO rescued with catalase from *H. pylori* demonstrated that excess CO_2_ release from photorespiration is suppressed with catalase activities far below native levels. These results indicate that once an activity threshold is met, decarboxylation reactions detrimental to plant growth are minimized, and any additional increase in catalase capacity may offer limited physiological benefit to enhancing *A*.

## Materials and methods

### Rescue catalase deficient lines (*cat2*-KO) with *H. pylori* isoforms

To express the *H. pylori* catalase isoforms in *A. thaliana*, the amino acid sequence of the katA catalase gene from *H. pylori* was identified in UniProt. This sequence was then codon optimized for higher plants, made compatible for Golden Gate cloning by removing the internal type II restriction sites, and targeted to the peroxisome by the addition of a type 1 peroxisome targeting signal (SKL) to the carboxyl terminus. Constructs were synthesized using a commercial service (Genewiz). The synthesized construct was then assembled into a Level 0 Golden Gate plasmid using a parts kit obtained from AddGene (Engler *et al*., 2014). Level 0 parts were then assembled into a Level 2 transformation vector under the rubisco small subunit promoter, an act2 terminator and kanamycin resistance marker (Supplemental Figure 1).

The *H. pylori* catalase isoform was transformed into *cat2*-KO *A. thaliana* lines by floral dipping for *in vivo* and *in vitro* functional analysis. *Cat2*-KO were used as the transformation background since these lines have minimal leaf expression of native catalase. Transformation were performed according to (van Hoof *et al*., 1996) with the following modifications. In brief, the binary vector was introduced into the *Agrobacterium tumefaciens* strain GV3101 (C58C1 Rifr) pMP90 by electroporation. Plants of ecotype Col-0 were grown under a photoperiod of 16h light /8 h dark at 20°C-22°C until the primary bolt was 5-15 cm long. A 500 mL culture of YEP medium containing kanamycin for selection was inoculated with *A. tumefaciens* and was resuspended in 1 L of infiltration medium. Arabidopsis plants were infiltrated with this suspension under 400 mm Hg vacuum for 5 minutes then return to growth chamber. Plants were grown for seed. *H. pylori* transformants were selected on kanamycin and basta plates. After rounds of generating homozygous lines, the genotypes were confirmed by PCR using forward primer (5 - AACCATCAAGTGCAGACCAGT - 3) and reverse primer (5 - CAAAGCCTCTAGGGTCCCTCAC - 3) (Supplemental Figure 2). Three lines were obtained, Hp143, Hp432, Hp615 lines, that showed elevated expression of catalase activity.

### Confirming localization in the peroxisome

To confirm that *H. pylori* catalase was targeted to the peroxisome, we determined the localization of the catalase isoforms to peroxisomes using confocal microscopy and transient expression. Co-localization of the florescent peroxisomal dye BODIPY with *H. pylori* catalase N-terminally tagged with M-Cherry in *Nicotiana benthamiana* protoplasts was imaged using confocal microscopy (Supplemental Figure 3).

### Plant material and growth conditions

*A. thaliana* seeds were cold stratified in for 4-6 days in deionized water in 2mL Eppendorf tubes before sowing. *A. thaliana* plants were sown in 0.7 or 1.5 L pots on Sure-Mix potting soil (Michigan Grower Products, Inc.). Plants were grown in growth chamber at a day/night temperature of 23°C/18°C, with light intensity ranging between 80-100 μmol photons m^−2^ s^−1^ with no humidity control. For Chlorophyll a fluorescence and growth analysis the photoperiod was set to 12/12, but for gas exchange and catalase activity assays the photoperiod was switched to 6/18 to develop larger leaves appropriate for gas exchange analysis. Plants were fertilized weekly with ½-strength Hoagland solution.

### Preparing crude protein extract and catalase enzyme assay

Crude protein extracts were prepared from the youngest, fully expanded leaves of *A. thaliana*. Two leaf punches (52 mm^2^) were taken from *A. thaliana* using a cork borer, immediately frozen in liquid N_2_, and stored at −80°C. Leaf punches were homogenized on ice with 0.5 mL of the Extraction buffer (50 mM EPPS buffer, pH 8.0, containing 1 mM EDTA, 10 mM DTT, 0.1% Triton X-100 [v/v], 0.5% polyvinylpyrrolidone, and 20 μL 1X SigmaFAST Protease Inhibitor Cocktail, EDTA Free (Sigma, St. Louis, MO, USA)), using a 2 mL glass-to-glass homogenizer (Kontes Glass Co., Vineland, NJ, USA). The homogenate was transferred into a 2 mL plastic Eppendorf tube and clarified by centrifugation for 15 min at 15,000 g and 4⍰C (Eppendorf Centrifuge 5424R, Eppendorf, Enfield, CT, USA). The supernatant, containing the clarified crude protein extract, was used for catalase activity assays.

The activity of catalase was determined in *A. thaliana* lines by the production of O_2_ (Aebi, 1983; Zelitch, 1989) using a Clark-type electrode with the following modifications. The reaction mix (50 mM K-phosphate buffer, pH 8.1) and crude protein extract were incubated for 30 sec to determine the O_2_ baseline. The reaction was initiated with 30 mM H_2_O_2_ and the increase in O_2_ production (nmol/mL) was observed for 1 min with 1 sec interval using a Oxygraph+ Oxygen Monitoring System (Hansatech Instruments Ltd). The initial rate of reaction was determined during the first 10 sec and the specific activity was expressed as μmoles O_2_ produced m^−2^ s^−1^ (Escobar *et al*., 1990; Szczepanczyk *et al.*, 2023).

### Quantifying CO_2_ release from a post illumination CO_2_ burst

Gas exchange was measured on the youngest, fully-expanded leaf using a LI-6800 (LI-COR Biosciences) with a 2cm^2^ chamber with 50:50 blue:red LEDs. Plants were measured according to (Gregory *et al*., 2024b). In brief, leaves were stabilized at 40 Pa CO_2_ at a leaf temperature of 25°C before starting, which usually took ∼20 minutes. They were then measured for a total of 30 minutes, initially under a light (10 minute; 1000 μmol PAR m^−2^ s^−1^), then dark (20 minutes; 0 μmol PAR m^−2^ s^−1^) period with logging every 1 seconds. Both steady-state assimilation (*A*_*s*_) and respiration in the dark (*R*_*D*_) were estimated from the last 10 points in the light and dark, respectively. A linear regression was fit using a baseline correction as the y-intercept, and a slope of 0. The baseline correction was identified during the last 200 seconds in the dark (Bao *et al*., 2021). The total amount of CO_2_ release during the PIB was estimated as the sum up the differences between the linear regression and the measured assimilation values within the burst (Gregory *et al*., 2024b).

### Estimating rates of *v*_*o*_

*v*_*o*_ for WT, *cat2*-KO, Hp143, Hp432, and Hp615 were estimated from measured gas exchange according to

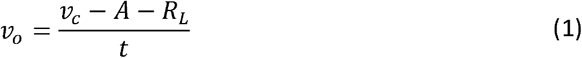

where *v*_*c*_ was determined by

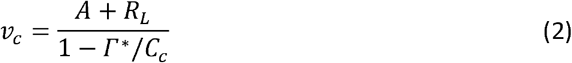

(Walker *et al*., 2020). The partial pressure of CO_2_ at the site of rubisco catalysis (*C*_*c*_) was determined by

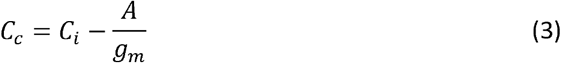

where *A* and *C*_*i*_ were measured at saturating light (1000 μmol PAR m^−2^ s^−1^) during the light phase of the PIB curve. *g*_*m*_, *Γ*, R*_*L*_, and t were previously measured or estimated in WT (2.23 μmol m^−2^ s^−1^ Pa^−1^, 4.482 Pa, 0.51 μmol m^−2^ s^−1^, 0.5) and *cat2*-KO mutants (2.01 μmol m^−2^ s^−1^ Pa^−1^, 5.827 Pa, 0.53 μmol m^−2^ s^−1^, 0.64) (Bao *et al*., 2021). Using the CO_2_ burst size as a proxy from the PIB measurement, the *H. pylori* mutants were classified with the same diffusional and biochemical parameters (*g*_*m*_, *Γ*, R*_*L*_, and *t*) as *cat2*-KO (i.e., Hp143) or WT (i.e., Hp432 and Hp615).

### Chlorophyll *a* fluorescence screening during an oxygen transient

For Chlorophyll *a* (Chl a) fluorescence analysis, a Dynamic Environment Photosynthetic Imager (Cruz *et al*., 2016) was coupled to a gas mixing system controlled by mass flow controllers (Alicat Scientific, Inc, USA) to monitor chlorophyll a fluorescence parameters during a transition from low to high photorespiratory conditions. Nitrogen (N_2_; Peak Scientific, USA), carbon dioxide (CO_2_; Airgas Specialty Gases, USA), and O_2_ (Airgas Specialty Gases, USA) were mixed and passed into an acrylic chamber (Plas-Labs, Lansing, MI, 63.5cm x 33cm x 17.8cm base with a 20° angle roof to minimize back-reflection of actinic and measuring light) to change the composition of the atmosphere surrounding plants during the screen, which took approximately 40 minutes to move from 2% to 40% O_2_.

Chl a fluorescence was imaged on an entire rosette of *A. thaliana* WT, *cat2*-KO, and three *H. pylori* transformed lines (Hp143, Hp432, and Hp615) to evaluate photosynthetic performance during an oxygen transient screen (Smith, 2024). The imaging protocol lasted three hours. Plants were first dark-adapted for one hour to capture the dark-adapted fluorescence yield (*F*_*o*_), and maximum quantum yield (*f*_*v*_*/f*_*m*_), the latter measured immediately following the dark adaptation. After this period, actinic lights (300 μmol PAR m^−2^ s^−1^) were turned on for the remaining two hours. During the three hours, a saturation flash was applied every two minutes to capture steady-state fluorescence (*f*_*s*_) and maximum fluorescence under light (*f*_*m*_⍰), enabling the calculation of parameters such as non-photochemical quenching (NPQ) and quantum yield of photosystem II (ϕ_II_) (Baker, 2008).

Simultaneously, plants were subjected to a gas-switching protocol designed to simulate a photorespiratory transition. For the first 1.6 hours, plants were exposed to low photorespiratory pressure (2% O_2_, 400 ppm CO_2_, balance N_2_). At 1.6 hours, the atmosphere was switched to high photorespiratory pressure (40% O_2_, 400 ppm CO_2_, balance N_2_) for the remaining 1.3 hours. All raw images were processed using the Visual Phenomics platform (Cruz *et al*., 2016).

### Measuring non steady-state *A* with Dynamics Assimilation Technique

Gas exchange was measured using the Dynamic Assimilation Technique (DAT) on the youngest, fully-expanded leaf of 8- to 9-week old plants using a LI-6800 (LI-COR Biosciences) with a 2cm^2^ chamber with 50:50 blue:red LEDs. Range matching and dynamic calculations were performed according to the manufacturer’s instructions and (Tejera-Nieves *et al*., 2024a; Tejera-Nieves *et al*., 2024b). Plants were measured under saturating light (1000 μmol PAR m^−2^ s^−1^) from 5 Pa CO_2_ to 150 Pa CO_2_ with a flow rate of 200 μmol s^−1^ at a leaf temperature of 25°C. Leaves were equilibrated at 40 Pa CO_2_ at the measuring temperature before starting a monotonic increase in CO_2_ starting at 5 Pa CO_2_.

CO_2_ response curves were fitted for the maximum rate of rubisco carboxylation (*v*_*c,max*_), and maximum rate of electron transport (*J*), and triose phosphate utilization (*TPU*; while a clear TPU limitation was not observed in all lines, we wanted to mitigate *J* overestimation). *v*_*c,max*_, *J*, and *TPU* (not reported) were estimated using R-based ACi fitting tool (Gregory *et al*., 2021; Saathoff *et al*., 2021) (see https://github.com/poales/msuRACiFit to access the R script with user-friendly interface).

### Plant growth and leaf trait measurements

Relative growth rate (RGR) was calculated in ImageJ from projected leaf area images taken every two days after germination (image collection was stopped once leaves overlapped and true area could not be obtained) (Schneider *et al*., 2012). Leaf number was counted a week prior before bolting, subsequently plant biomass was cut above ground and weighed to get fresh mass (g). During the biomass harvest plants were imaged, then dissected to measure projected leaf area in ImageJ (m^2^). Leaves were then placed in an envelope and dried at 60°C for 48 hours and weighed for leaf dry mass (g). Leaf areas and dried weights were used to calculate dry leaf mass per area (LMA; g m^−2^) and specific leaf area (SLA; m^−2^ kg). Fresh and dried masses were used to resolve water content in tissue on a percentage basis.

### Data processing and statistical analysis

Growth analysis, Chla fluorescence, gas exchange and biochemical data were analyzed and visualized using custom scripts in R (R Core Team, 2021). We used emmeans() in the emmeans package for mean and parameter comparison. Growth analysis, gas-exchange and biochemical data were checked for a normal distribution (assumption #1) using the shapiro.test() in stats() package and for homogeny of variance (assumption #2) using the leveneTest() in car() package before proceeding with parametric testing. To check for statistical differences between *cat2*-KO and the three heterologous *H. pylori* lines (all within the same genetic background), growth analysis, gas exchange and biochemical data were analyzed using pairwise.t.test() from the stats() package with Bonferroni correction, to control for Type I errors. Growth analysis, gas exchange and biochemical data were visualized using geom_boxplot() to represent the distribution of the variable of interest within a genotype. The box represents the first and third quartile, and the line inside of the box represents the median. The whiskers represent the minimum and maximum values, while the solid dots represent outliers (calculated as 1.5* interquartile range); however, these were not removed from the visualization or analysis. All ANOVA tests were followed with a Tukey’s post-hoc test.

## Results

### Catalase capacity, photorespiratory influx, and their catalytic balance

To determine catalase capacity in *A. thaliana* lines and knockouts rescued with heterologous expression of catalase from *H. pylori*, we measured catalase activities under saturating substrate concentration to determine an *in vitro V*_*max*_ (Figure 1A). Catalase activities were measured in leaves of *A. thaliana* at 25°C using crude protein extract (Supplemental Table 1). Cat2-KO had 6% of the total catalase activity as WT lines which aligns well with 10% that been measured previously (Bao *et al*., 2021; Mhamdi *et al*., 2010). When comparing the *H. pylori* catalase expressing lines to *cat2*-KO, which is in the genetic background, the mean activities increased by 1.62 (Hp143), 1.83 (Hp432), 2.86 (Hp615) fold. However, only the Hp615 heterologous expressing line had a significant difference (*p* < 0.05) in total catalase activity when compared to *cat2*-KO.

**Figure 1.**
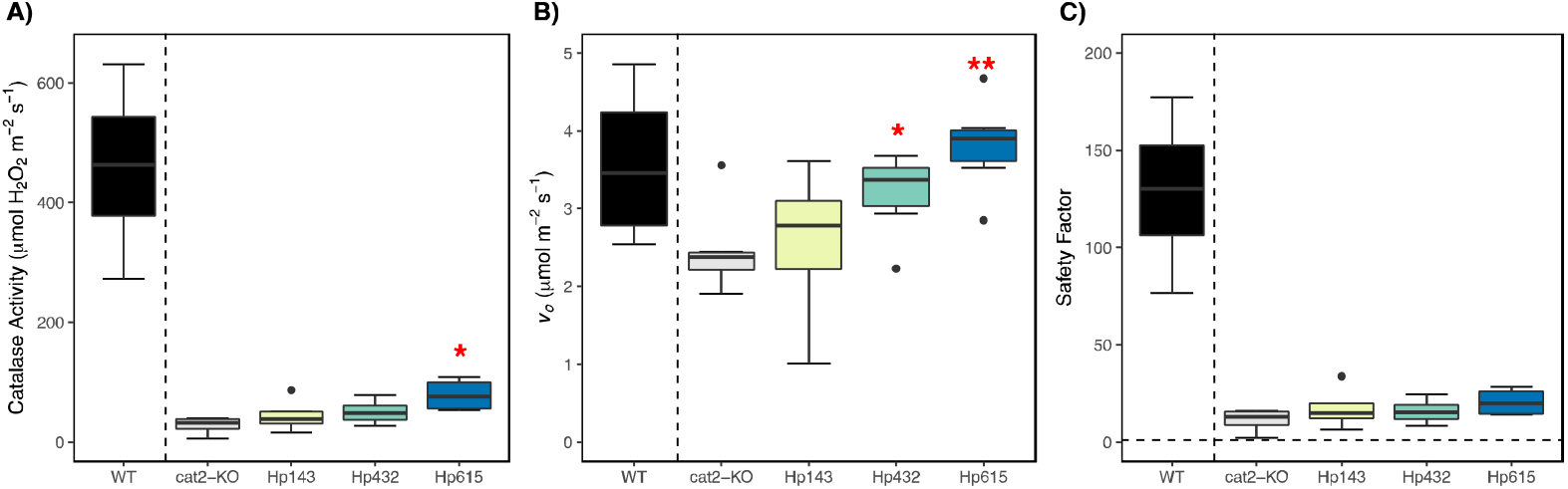
Catalase Activity, Photorespiratory Influx, and Safety Factor in *Arabidopsis thaliana*. A) Specific activities per m^2^ leaf area were measured in *A. thaliana* at 25°C using crude protein extract for catalase, B) photorespiratory influx (*v*_*o*_) were estimated in *A. thaliana* at 1000 μmol PAR m^−2^ s^−1^ and 25°C using a LI-COR LI-6800 infrared gas analyzer, and C) safety factors were calculated by dividing catalase activity by *v*_*o*_ for each genotype. Colors represent genotype. Shown are the boxplots indicating the biological replicates (n = 4-6). Significant difference between the *cat2*-KO and individual heterologous lines are indicated by one (*p* ≤ 0.05), two (*p* ≤ 0.01), or three (*p* ≤ 0.001) red asterisk as determined by pairwise t-test with Bonferroni correction.

Since the substrate for catalase, H_2_O_2_, is produced in a 1:1 ratio with the velocity of rubisco oxygenation (*v*_*o*_), quantifying *v*_*o*_ will estimate rates H_2_O_2_ influx in the peroxisome. To assess rates of photorespiratory influx from *v*_*o*_, we estimated the *v*_*o*_ and the velocity of rubisco oxygenation per carboxylation (*v*_*o*_*/v*_*c*_) using steady-state assimilation data at saturating light (1000 μmol PAR m^−2^ s^−1^) from PIB curves. *Cat2*-KO had significantly lower *v*_*o*_ and *v*_*o*_ */v*_*c*_ when compared to Hp432 (*p* < 0.05 and *p* < 0.05) and Hp615 (*p* < 0.05 and *p* < 0.001) lines but had similar rates to Hp143 at 25°C (Figure 1B & Supplemental Table 1).

To evaluate the relationship between photorespiratory influx and downstream catalase capacity to enzymatically process H_2_O_2_, safety factors were calculated to quantify the excess capacity (Figure 1C and Supplemental Table 1) (Alexander, 1981; Diamond, 2002; Gregory *et al*., 2024a). The safety factor metric helps reveal the balance between influx and capacity and can resolve whether enzymes activities fall short of *v*_*o*_ (safety factors < 1), are quantitatively matched with *v*_*o*_ (safety factors = 1) or maintain excess capacity to process *v*_*o*_ (safety factors > 1). To calculate the safety factor, catalase *V*_*max*_ measured in leaves at 25°C was divided by *v*_*o*_ estimated at 25°C. In the *A. thaliana* lines measured, safety factors were greater than 1 in WT, *cat2*-KO, Hp143, Hp432 and Hp615 lines, indicating that each line maintains a more than enough enzymatic capacity to process H_2_O_2_ produced following rubisco oxygenation. However, WT had a significantly larger safety factor than *cat2*-KO, Hp143, Hp432, and Hp615. Although there are slight differences in *v*_*o*_ between the lines, the lower safety factors are driven mainly by the low absolute catalase activities of *cat2*-KO (27.5 μmol H_2_O_2_ m^−2^ s^−1^), Hp143 (44.7 μmol H_2_O_2_ m^−2^ s^−1^), Hp432 (50.4 μmol H_2_O_2_ m^−2^ s^−1^), and Hp615 (78.7 μmol H_2_O_2_ m^−2^ s^−1^) compared to WT (457.7 μmol H_2_O_2_ m^−2^ s^−1^). We next wanted to determine if these catalase activities in lines expressing *H. pylori* catalase, while well below WT levels, were sufficient to rescue other aspects of *Cat2*-KO physiology.

### Evidence for excess CO_2_ release rescue from Post Illumination CO_2_ Burst

To determine whether the excess photorespiratory CO_2_ release from photorespiration was rescued in Hp143, Hp432, and Hp615, we measured the Post Illumination CO_2_ Burst (PIB; Figure 2). The PIB was quantified for the CO_2_ burst (i.e., integrated area under the PIB) and steady-state assimilation (i.e., averaged of the last 10 points in the light) using an R-based fitting tool (Gregory *et al*., 2024b) (Figure 2B & C). Our measurements of the PIB reveal the CO_2_ burst was greater in *cat2*-KO when compared to WT lines, which agrees with photorespiratory CO_2_ evolution found previously in (Bao *et al*., 2021; Keech *et al*., 2012). Hp143 had a CO_2_ burst size similar to *cat2*-KO, while Hp432 and Hp615 matched WT. Steady-state *A* was lower in *cat2*-KO and Hp143 compared to WT, Hp432, and Hp615 lines. This decrease in steady-state *A* supports the explanation that CO_2_ evolution from photorespiration increases through NED or other CO_2_-releasing reactions in *cat2*-KO and Hp143. Additionally, *g*_*sw*_ remained statistically similar in the WT, *cat2*-KO, Hp143, Hp432, and Hp615 lines, indicating that the stomata did not differentially limit CO_2_ diffusion between lines (Supplemental Table 1). However, *C*_*i*_ was lower in Hp615 compared to *cat2*-KO, Hp143, Hp432, and WT lines, which corresponds to the steady-state A being the highest (Supplemental Table 1).

**Figure 2.**
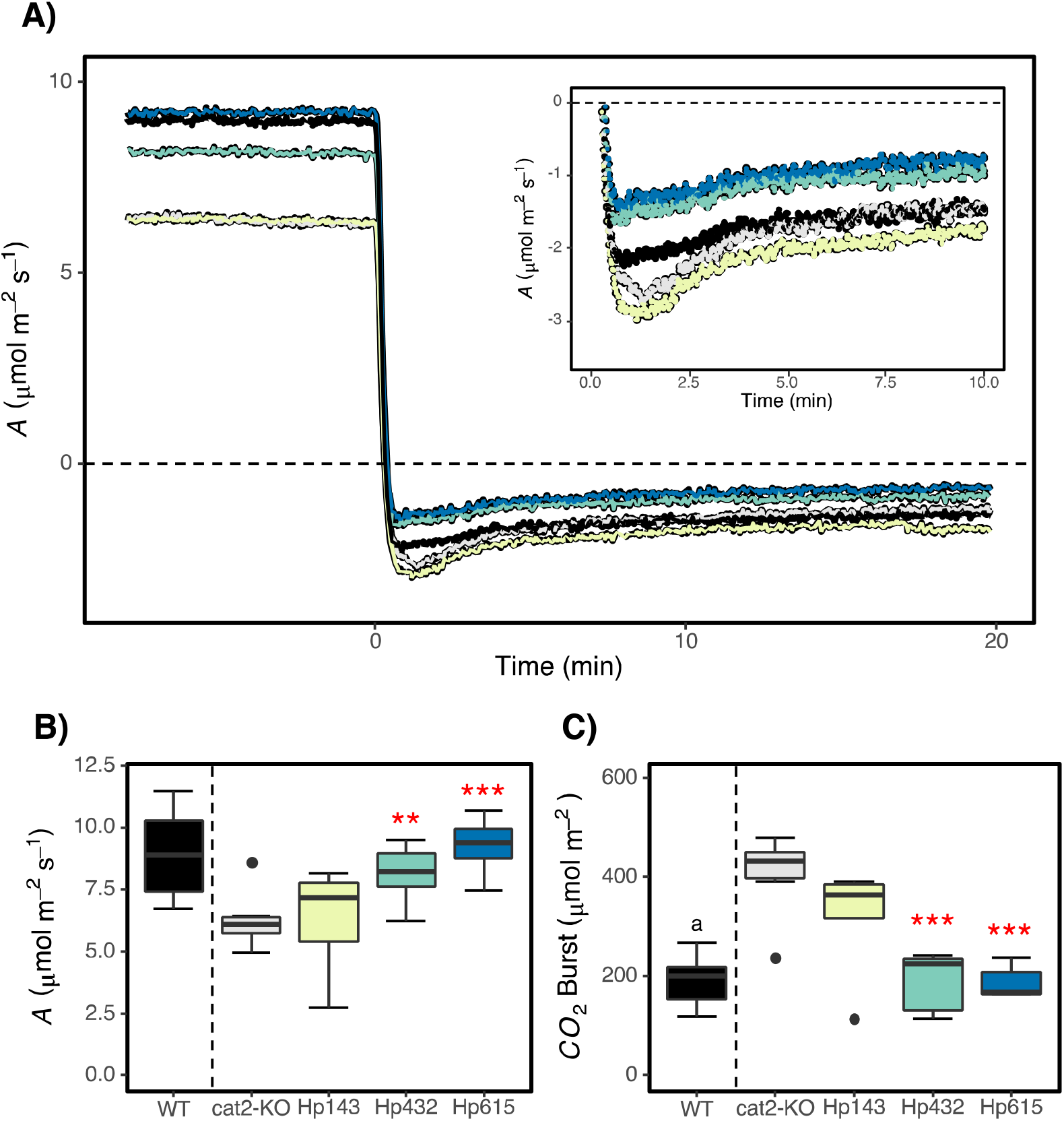
Post Illumination CO_2_ Burst curves at 25°C in *Arabidopsis thaliana*. A) Net carbon fixation (*A*) was measured in *A. thaliana* during a light (1000 μmol PAR m^−2^ s^−1^) to dark transient for 30 minutes using a LI-COR LI-6800 infrared gas analyzer, B) Steady-state *A* was averaged across the last 10 seconds in the light, and C) CO_2_ burst size was quantified by integration of the peak area of CO_2_ release. Colors represent genotype. Shown are the boxplots indicating the biological replicates (n = 6). Significant difference between the *cat2*-KO and individual heterologous lines are indicated by one (*p* ≤ 0.05), two (*p* ≤ 0.01), or three (*p* ≤ 0.001) red asterisk as determined by pairwise t-test with Bonferroni correction.

### Evaluating CO_2_ response curves and the biochemical limitations of photosynthesis

To understand how this reduced carbon loss from photorespiration impacts net gas exchange, we measured CO_2_ response curves using a dynamic assimilation technique. CO_2_ response curves were measured in each genotype and *v*_*c,max*_ and *J* were estimated at 25°C in WT, *cat2*-KO, Hp143, Hp432, and Hp615 (Supplemental Table 2 and Figure 4).

**Figure 3.**
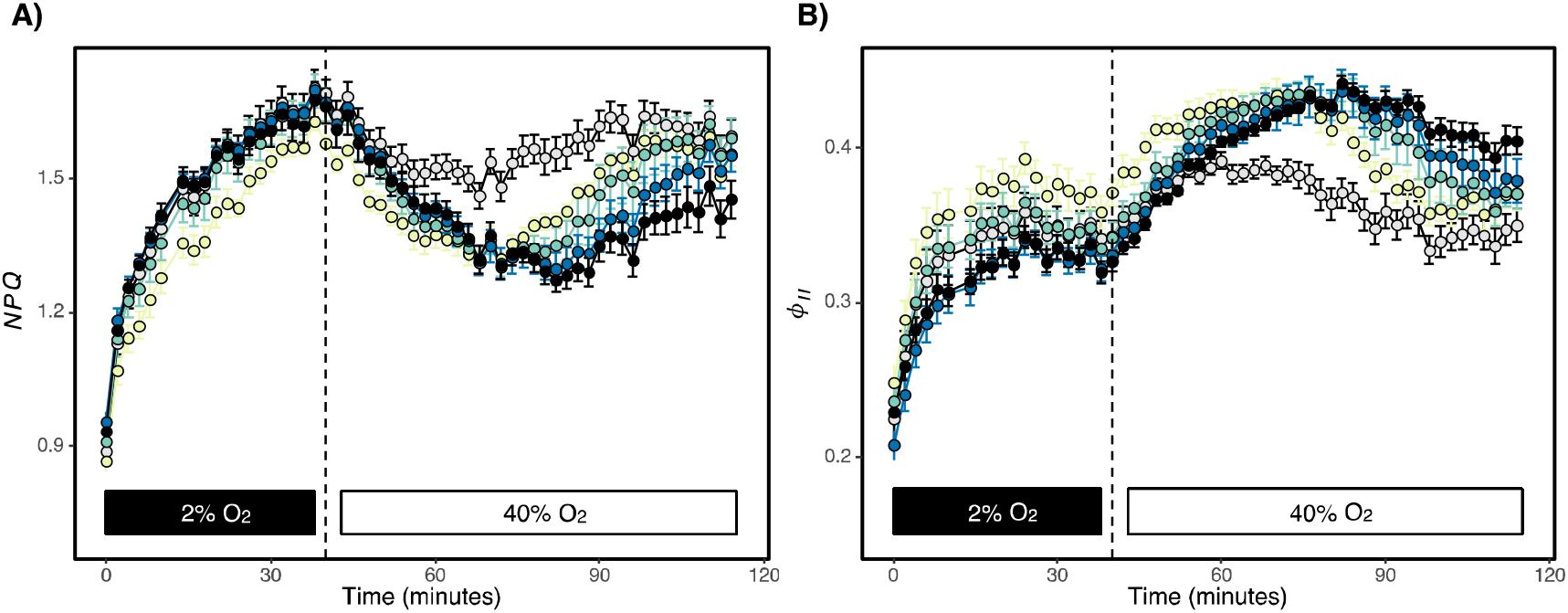
Chlorophyll fluorescence screen across an oxygen transient at 25°C in *Arabidopsis thaliana*. A) NPQ and B) *ϕ*_*II*_ were screened in *A. thaliana* for 2 hours in the light (300 μmol photons m^−2^ s^−1^) using Dynamic Environmental Photosynthetic Imager (DEPI). Colors represent genotype, where WT (black), *cat2*-KO (grey), Hp143 (yellow), Hp432 (green), and Hp615 (blue). Shown are the mean of the biological replicates (n = 8) with standard error bars.

**Figure 4.**
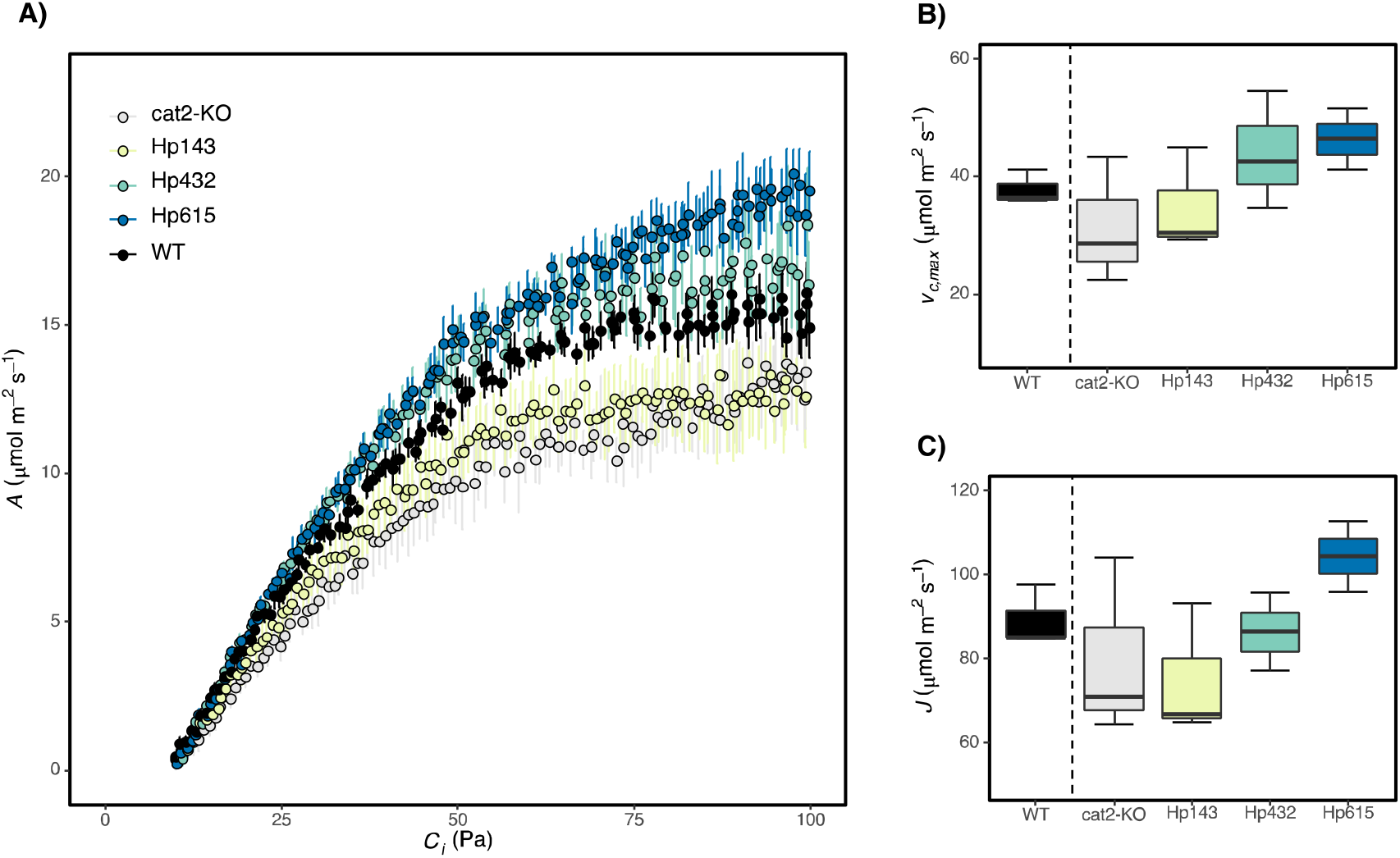
CO_2_ response curves at 25°C in *Arabidopsis thaliana*. A) Net carbon fixation (*A*) was measured in *A. thaliana* during a monotonic increasing CO_2_ response curve at 1000 μmol PAR m^−2^ s^−1^ using a LI-COR LI-6800 infrared gas analyzer, B) *v*_*c,max*_ estimation, C) *J* estimation between genotypes. Colors represent genotype. Shown are the boxplots indicating the biological replicates (n = 3). No significant differences were determined between *cat2*-KO and individual heterologous lines for *v*_*c,max*_ or *J*.

*v*_*c,max*_, and J were similar across the genotypes (Supplemental Table 1). Notably, the mean comparisons of *v*_*c,max*_, and J reflect the catalase activity in Hp143, Hp432, and Hp615, where the lowest activity has the lowest parameter estimates (i.e., Hp143) and the highest activity has the highest parameter estimates (i.e., Hp615).

A was compared under *C*_*i*_ values that were rubisco-limited (12-15 Pa & 32-34 Pa), J-limited (32-34 Pa, 52-55 Pa, 72-75 Pa, 92-95 Pa), and TPU-limited (92-95 Pa, although a clear TPU limitation was not observed in all lines). Under rubisco-limited conditions WT, *cat2*-KO, Hp143, Hp432 and Hp615 all had similar rates of *A*. Under *J*-limited conditions, WT, Hp432, and Hp615 had greater rates of A than *cat2*-KO and Hp143. Under TPU-limited conditions, Hp432 and Hp615 had greater rates of *A* than WT, *cat2*-KO, and Hp143 (Supplemental Table 2).

### Chlorophyll a (*Chl a*) fluorescence screening under O_2_ transient reveals an NPQ phenotype

To evaluate the influence of catalase activity on the light reactions of photosynthesis. we measured non-photochemical quenching (NPQ) under a 2% to 40% oxygen transient in WT, *cat2*-KO, Hp143, Hp432, and Hp615 lines (Figure 3). The induction of NPQ under 40% (high photorespiratory pressure) is a well-characterized fluorescence phenotype of photorespiratory mutants (Smith, 2024). Under 2% O_2_, WT, cat2-KO, and the three independent *H. pylori* lines had similar NPQ. However, once the atmosphere switched to 40% O_2_, the NPQ diverged between WT and *cat2*-KO after ∼20 minutes. WT had a lower NPQ, while *cat2*-KO maintained a higher NPQ for the rest of the measurement. Hp143 diverged from WT first, followed by Hp432, then Hp615. The difference between the NPQ of *cat2*-KO and Hp615 under 40% oxygen correlates with the greater catalase activity.

To assess the influence of catalase activity on productive photochemistry, the quantum yield of photosystem II (ϕ_II_) was also measured during the 2% - 40% O_2_ transient. Consistent with the NPQ results, as the atmosphere shifted to 40% O_2_ and NPQ adjusted to high (*cat2*-KO) or low (WT, Hp143, Hp432, Hp615) levels, ϕ_II_ inversely adjusted.

### Assessing Plant Growth

To determine if the catalase activities in the heterologous *H. pylori* lines is sufficient to rescue the excess CO_2_ release and stunted growth phenotype of *cat2*-KO lines, we measured various growth metrics (Supplemental Table 1). Relative growth rate (RGR) provides information on the plants’ overall growth rate per unit time and indicates efficient resource acquisition and utilization. Dry leaf mass per area (LMA) is a complex variable that provides information on leaf structural properties, mainly leaf thickness and density. Our measurements reveal that Hp615 and Hp432 rescued the leaf number, RGR, and LMA relative to *cat2*-KO (Figure 5B, 5C, 5D). For the Hp615 line, the trend of increased plant growth relative to *cat2*-KO correspond with the increased catalase activity (Figure 1A; *p* < 0.05).

**Figure 5.**
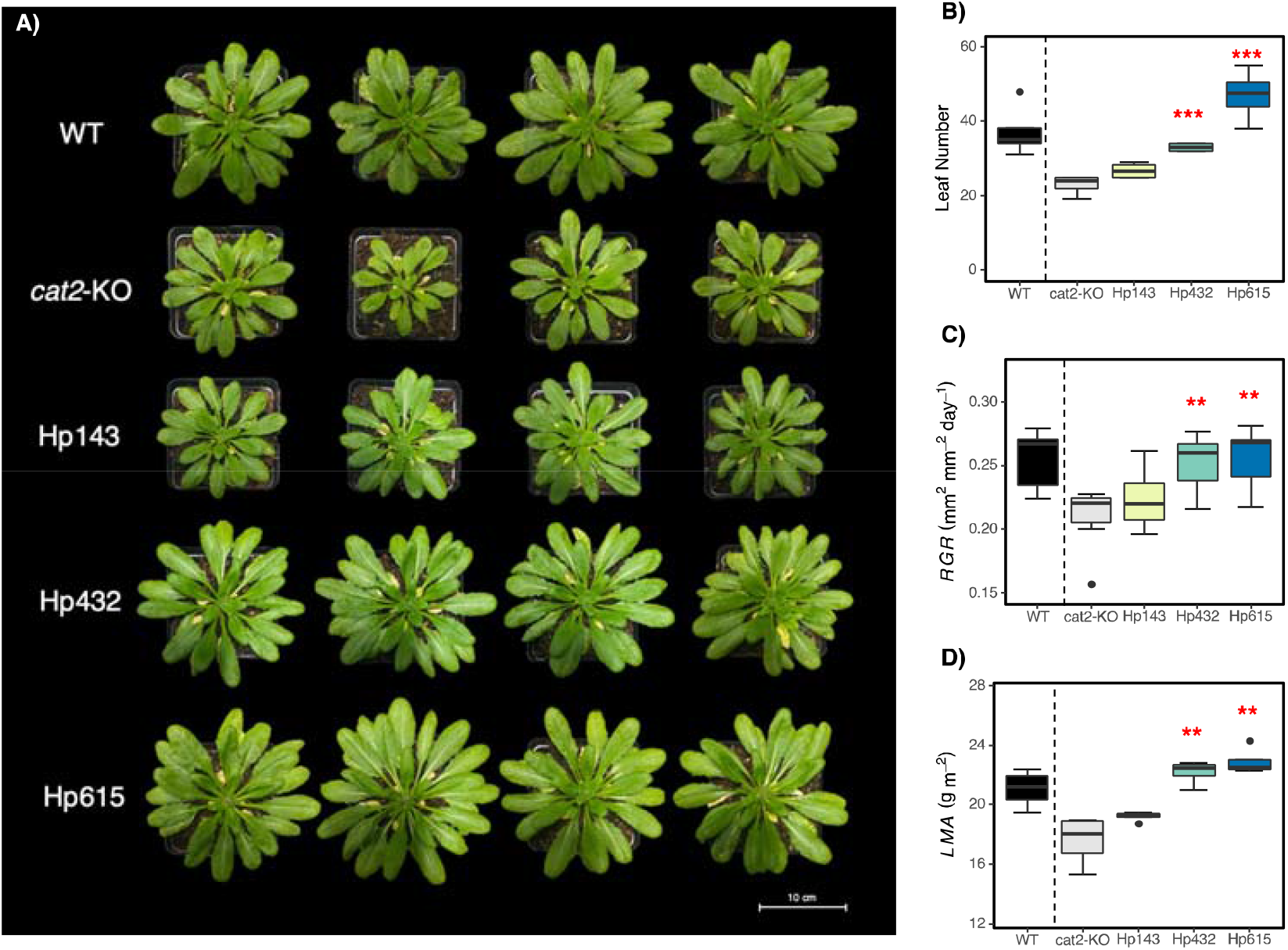
Growth Analysis of *Arabidopsis thaliana* genotypes. A) Four replicates of each genotype were grown under 23°C/18°C and a 12/12 photoperiod at 80-100 μmol photons m^−2^ s^−1^, B) Leaf Number was counted a week before bolting, C) relative growth rate (RGR), and D) Leaf Dry Mass per Area was counted a week before bolting. Colors represent genotype. Shown are the boxplots indicating the biological replicates (n = 4-7). Significant difference between the *cat2*-KO and individual heterologous lines are indicated by one (*p* ≤ 0.05), two (*p* ≤ 0.01), or three (*p* ≤ 0.001) red asterisk as determined by pairwise t-test with Bonferroni correction.

## Discussion

### Efficient H_2_O_2_ degradation may not require excess capacity of catalase

This work revealed that the catalase activity in WT plants far exceeds what is needed to prevent excess CO_2_ loss from decarboxylation reactions and NPQ increases (Figure 1A). Lines with substantially lower catalase safety factors – such as 20.7 (Hp615) and 15.8 (Hp432) – exhibit no additional CO_2_ loss compared to WT (128.6), suggesting that photorespiratory function can be maintained even with a large reduction in WT catalase capacity under the experimental conditions measured in this study. Although the safety factors in Hp432 and Hp615 are not significantly higher than the *cat2*-KO, which exhibits clear physiological defects, the absence of a growth or photoprotective penalty in these lines suggests that they are still operating above a critical threshold for effective H_2_O_2_ scavenging. In other words, WT plants maintain 6- to 8-fold greater catalase capacity than what is strictly necessary to prevent deleterious accumulation of H_2_O_2_ compared to the *H. pylori* isoform-expressing lines.

This raises the question: why maintain such excess capacity? One possibility is that WT catalase levels provide a buffer against sudden or extreme oxidative stress conditions. Catalase catalyzes the dismutation of hydrogen peroxide, using two H_2_O_2_ molecules—one as an oxidant and one as a reductant—to form water and oxygen. This mechanism, while energetically favorable, also results in a high *k*_*m*_, meaning catalase only operates efficiently at high substrate concentrations. Therefore, high enzyme abundance is required to maintain effective H_2_O_2_ detoxification, especially under transient spikes in H_2_O_2_ production. Unlike other peroxidases, such as ascorbate peroxidase and glutathione/thioredoxin peroxidase, catalase does not rely on co-substrates or reducing equivalents and is thus metabolically less costly (Mhamdi *et al*., 2010; Willekens *et al*., 1995). This independence from cellular redox networks may make catalase particularly valuable for rapid, bulk detoxification of H_2_O_2_ during high-flux events and additionally explain why excess investment in catalase is not detrimental.

Indeed, certain physiological scenarios, such as when stomata are closed mid-day under high light and/or high temperature, can induce exceptionally high rates of photorespiration. In these conditions, starch degradation may drive an increase in triose phosphate concentrations in the chloroplast, which can fuel the regeneration of RuBP and sustain rubisco activity, leading to high rates of glycolate and H_2_O_2_ production through photorespiration. Under the substantial rates of H_2_O_2_ production in this scenario, WT catalase activity may be pushed closer to its physiological limits, potentially explaining the apparent excess capacity under moderate conditions tested in this study. This excess capacity may additionally reflect strong evolutionary pressure to buffer against short, transient oxidative bursts, where insufficient scavenging could cause costly cellular damage. In this context, maintaining a high safety factor for catalase likely serves as a preventative strategy, minimizing the risk of cellular injury during unpredictable H_2_O_2_ spikes.

### Peroxisomal catalase plays an essential role in maintaining plant growth

This study demonstrates that peroxisomal catalase plays an essential role in plant growth by preventing excess CO_2_ loss and reducing the impact of H_2_O_2_ on the light reactions. The photorespiratory mutant, *cat2*-KO, which only has 6% of WT catalase activity, exhibited a severe growth phenotype linked to greater CO_2_ release and higher NPQ due to insufficient degradation of H_2_O_2_ (Figure 1A, Figure 2C, Figure 3A). Although both WT and *cat2*-KO show sufficient catalase capacity according to *in vitro* assays under saturating substrate concentrations (30 mM H_2_O_2_; safety factor: WT = 128.6 and *cat2*-KO = 11.1), this capacity likely overestimates *in vivo* performance. Most enzymes, including catalase, do not operate at their *V*_*max*_ in vivo but instead operate closer to their *K*_*m*_ (*K*_*m*_ = ½ *V*_*max*_) due to a lower concentration of substrate and the need to maintain catalytic efficiency (Cornish-Bowden, 1976; Somero, 1978). For example, H_2_O_2_ concentrations in the peroxisome of *A. thaliana* have been estimated at 10 mM (Foyer *et al*., 2016), which would result in reduced in vivo catalase activity in both lines. Under these physiological conditions, *cat2*-KO likely cannot meet the metabolic demand for H_2_O_2_ detoxification at WT rates of *v*_*o*_, contributing to the severe growth inhibition.

The growth phenotype and decrease in A in *cat2*-KO lines is in part explained by the increase in photorespiratory CO_2_ release compared to WT lines. *Cat2*-KO maintains a lower steady-state and non-steady-state A than WT lines measured in both the PIB and CO_2_ response curve at 25°C under ambient CO_2_ concentrations, respectively (Figure 2B and Supplemental Table 2). The decrease in A was not associated with CO_2_ diffusion limitation through the stomata, since *g*_*sw*_ was similar across WT and *cat2*-KO lines (Supplemental Table 1). However, the excess CO_2_ release in *cat2*-KO evaluated using the CO_2_ burst, accounts for the loss in *A* (Figure 2B). This CO_2_ burst, measured during a light-dark transient, is an indication of the photorespiratory release of CO_2_ under steady-state under illumination. The additional CO_2_ release due to the H_2_O_2_ accumulation is hypothesized to come from multiple sources. The sources include both the increased frequency of NED reactions and CO_2_ release from the G6P shunt (Bao *et al*., 2021; Li *et al*., 2019b; Sharkey *et al*., 2016). *Cat2*-KO likely stimulates the G6P shunt if H_2_O_2_ accumulation causes 2-PG buildup. A recent report shows that accumulation of H_2_O_2_ promotes the sulfenylation of phosphoglycolate phosphatase and inhibits its activity (Fu *et al*., 2024). The reduction in activity would promote the accumulation of 2-PG, which would inhibit triose phosphate isomerase, and slow the C_3_ cycle. This inhibition of metabolism could be overcome with the activation of the G6P shunt (Li *et al*., 2019b). Therefore, the excess CO_2_ burst in *cat2*-KO lines would agree with the contribution of CO_2_ release from the G6P shunt as well as NED.

In addition to carbon metabolism, the accumulation of H_2_O_2_ directly influences the light reactions. H_2_O_2_ can be mobile within the cell and has the ability to travel across peroxisomal membranes or organelles through diffusive paths or mediated by specialized or general transporters (Dynowski *et al*., 2008; Mubarakshina *et al*., 2010; Visser *et al*., 2007). Additionally, there is evidence that once H_2_O_2_ accumulates in the chloroplast, it can activate cyclic electron flow (CEF) through preexisting CEF machinery (Strand *et al*., 2015). In the *Chla* fluorescence analysis, NPQ increases in *cat2*-KO after ∼20 minutes under high photorespiratory pressure (40% O_2_), and it is maintained at greater level compared to WT (Figure 3A). *Cat2*-KO maintains a higher NPQ because of the acidification of the lumen through CEF being activated by H_2_O_2_ (Li *et al*., 2019a; Strand *et al*., 2015). Although CEF was not measured in this study, an increase in CEF in *cat2*-KO would support the hypothesis that G6P shunt involvement, as excess ATP would need to be generated to energetically support the shunt (Li *et al*., 2019b).

Moreover, the increase in NPQ correlates with a decrease in ϕ_II_, which suggests a decrease in linear electron flow (LEF; Figure 3B). The decrease in LEF in *cat2*-KO reduces the amount of ATP and NADPH that is produced at a given light intensity, effectively reducing *A* under high photorespiratory pressure by limiting the energy available for the C_3_ cycle. Consistent with these findings, under *J*-limited conditions on the CO_2_ response curve, rates of *A* were lower in *cat2*-KO compared to WT (Figure 4 and Supplemental Table 2). This suggests that processes involved in RuBP regeneration—such as electron transport through the cytochrome b_6_f complex, which is required for both LEF and CEF—are limiting *A* in *cat2*-KO compared to WT (Busch *et al*., 2024; Johnson *et al*., 2021; Sharkey *et al*., 2007). Consistent with electron transport limiting *A*, and not a general decrease in photosynthetic capacity, under rubisco-limited conditions, *cat2*-KO had similar *A* rates as WT, indicating that the limitation in *A* occurs specifically during RuBP regeneration (Figure 4 and Supplemental Table 2).

These metabolic and photochemical disruptions significantly impaired growth in *cat2*-KO compared to WT, as reflected in reductions in leaf number, RGR, and LMA (Figures 5A–D). Together, our findings show that catalase deficiency disrupts both carbon fixation and energy metabolism via H_2_O_2_ accumulation, leading to reduced growth.

### The *H. pylori* catalase isoform confers a benefit to photosynthetic performance, with less activity

Two out of the three independent expression lines of *H. pylori* catalase rescued the *cat2*-KO growth phenotype back to WT growth, with less total catalase activity as compared to WT (Figure 1A, and Figure 5). The catalase activities for the one of the independent *H. pylori* catalase isoform expression lines (Hp615) was greater than the *cat2*-KO line. However, the other *H. pylori* lines (Hp143 and Hp432) were not statistically different in their activities, despite both expressing the *H. pylori* catalase (Supplemental Figure 2). Interestingly, *in vivo* estimates of the PIB from Hp432 and Hp615 were similar to WT while having 9 and 6-fold less catalase activity than WT, respectively (Figure 2B). Hp143, the expression line with the least catalase activity (10 less than WT), has a comparable PIB as the *cat2*-KO (Figure 2C). The reduction in CO_2_ burst from Hp432 and Hp615 reflected an increase in steady-state A, which was greater than *cat2*-KO. The gain in *A* to WT levels was due to the reduction in CO_2_ loss in the CO_2_ burst, rather than CO_2_ diffusion limitation differences caused by *g*_*sw*_, which were similar in all lines at steady-state under illumination (Figure 2 & 4 and Supplemental Table 1).

In addition, Hp432 and Hp615 broadly mirror the NPQ behavior of WT line under high photorespiratory pressure, which effectively increases *A* by increasing the amount of ATP and NADPH produced at a given light intensity compared to *cat2*-KO for the C_3_ cycle (Figure 3). In agreement with these findings, *A* rates under *J*-limited conditions were greater in Hp432 and Hp615 than *cat2*-KO. Interestingly, Hp432 and Hp615 rates of A were similar to WT at ambient conditions, which align with steady-state A from the PIB, but larger at moderate (72-75 Pa) and high (92-95 Pa) CO_2_ concentrations. This difference is attributed to WT lines becoming *TPU*-limited at high CO_2_ concentrations, while Hp432 and Hp615 remain *J*-limited. In addition, under *rubisco*-limited conditions on the CO_2_ response curve, Hp432 and Hp615 had similar *A* rates as *cat2*-KO, Hp143, and WT lines, revealing that the difference in A occurs during RuBP regeneration (Figure 4 and Supplemental Table 1).

Hp432 and Hp615 increase photorespiratory efficiency and prevent high NPQ which rescued the severe growth phenotype in *cat2*-KO back to WT growth under the experimental growth conditions (Figure 5A). The mean leaf number, RGR, and LMA in Hp432 and Hp615 increased by 1.4, 1.2, 1.3 and 2.0, 1.2, 1.3 compared to *cat2*-KO, respectively (Figure 5B, C, D). Hp432 and Hp615 had statistically similar leaf number, RGR and LMA to WT. The upward trend in these growth metrics between the three independent expression lines of *H. pylori* catalase reflect their catalase activity. The differences in catalase activities in Hp143, Hp432, and Hp615 are likely driven by expression differences through gene stability, or chromosomal positional effects (Bandopadhyay *et al*., 2010; Betts *et al*., 2019; Matzke *et al*., 1998; Strauss *et al*., 2016). Taken altogether, Hp432 and Hp615 have enough catalase capacity to rescue the cat2-KO growth phenotype to WT growth, but Hp143 does not.

## Acknowledgements

We would like to thank Cody Keilen and Nick Deason (MSU Growth Chamber Facility) for overall greenhouse maintenance and pest management during plant cultivation. We thank Kalia Smith for initially setting up the NPQ screen and helpful discussion regarding the chlorophyll fluorescence parameters.

## Data Availability

The data that supports the findings of this study are available from the corresponding author upon request.

## Supplemental Figures

**Supplemental Figure 1.**
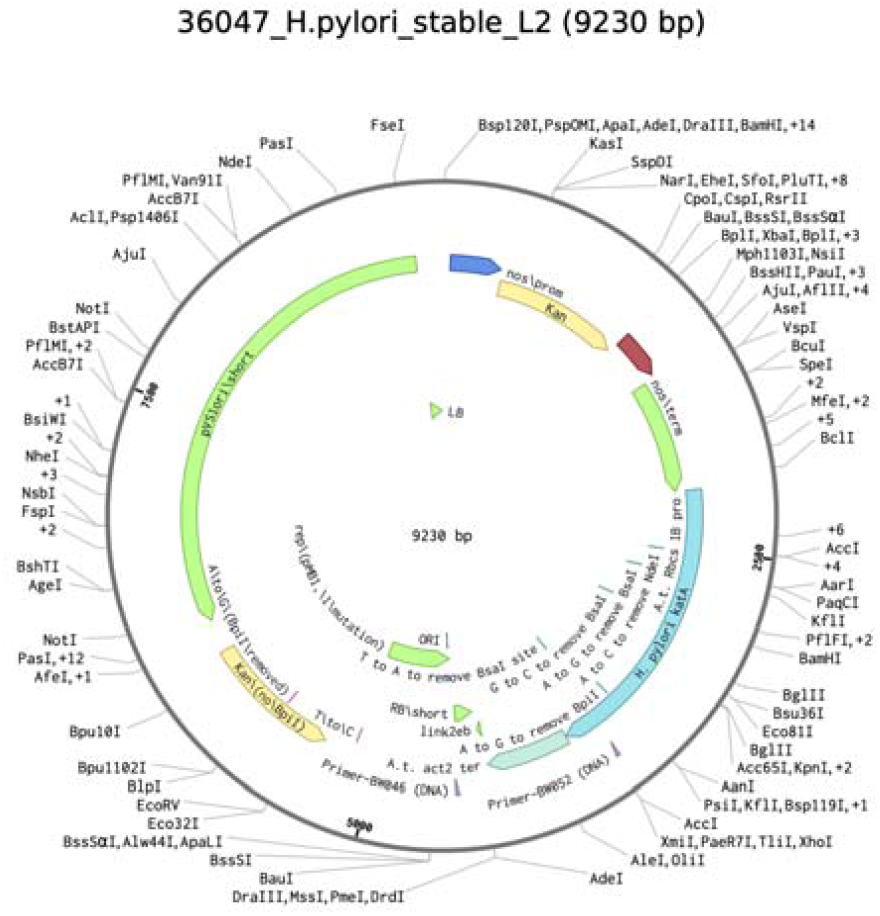
*Heliobacter pylori* transformation vector map.

**Supplemental Figure 2.**
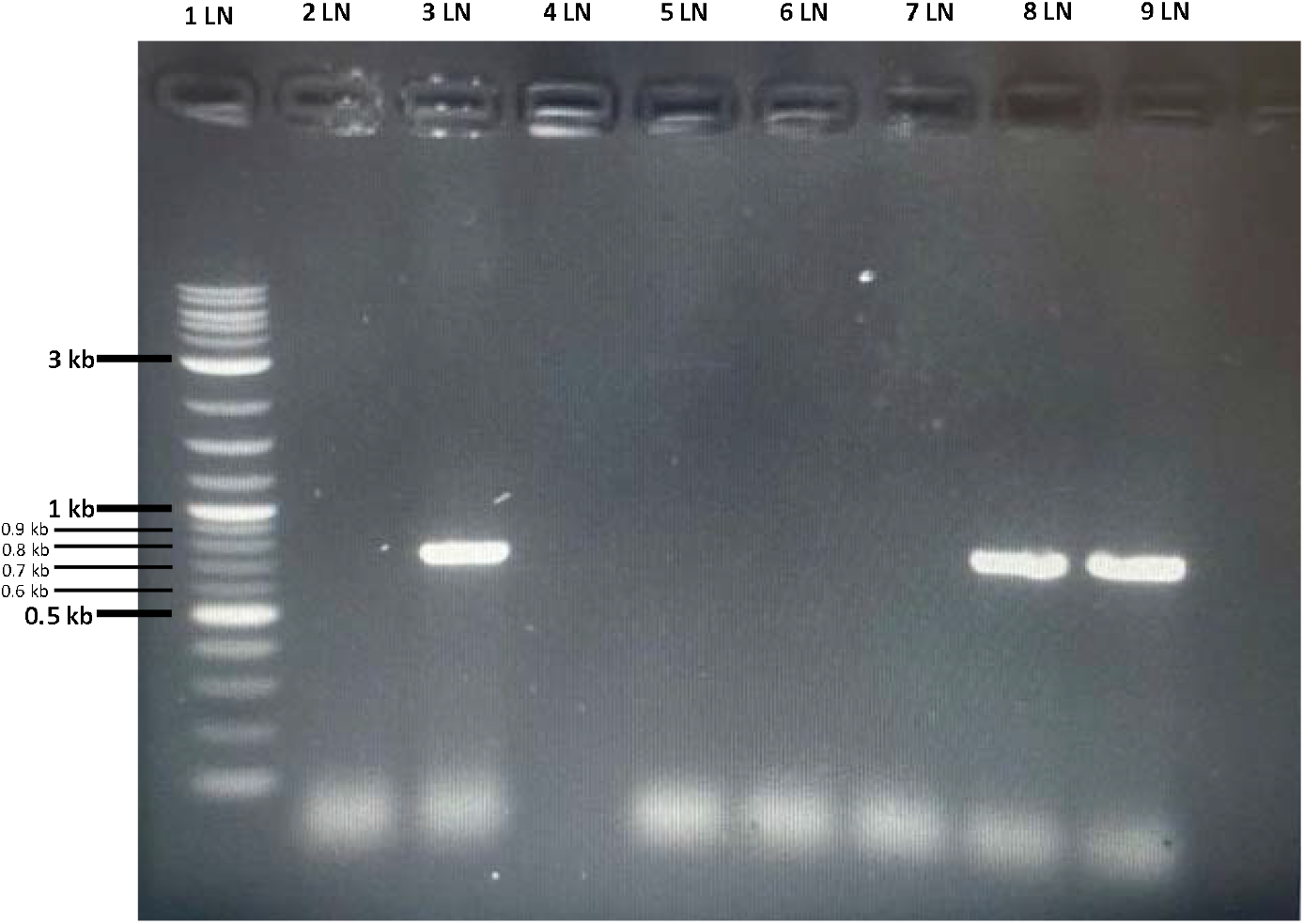
PCR genotyping in *Arabidopsis thaliana*. Agarose gel electrophoresis (1%) of PCR products used for genotyping various *Arabidopsis thaliana* lines. Lane 1: 1 kb Plus DNA ladder (ThermoFisher); Lane 2: *cat2*-KO knockout; Lane 3: Hp143; Lane 4: empty; Lane 5: wild type; Lane 6: *cat2*-KO knockout; Lane 7: Hp143 (low-quality DNA extraction, no band visible); Lane 8: Hp432; Lane 9: Hp615. PCR was performed using the following oligonucleotide primers: forward 5′-AACCATCAAGTGCAGACCAGT-3′ and reverse 5′-CAAAGCCTCTAGGGTCCCTCAC-3′.

**Supplemental Figure 3.**
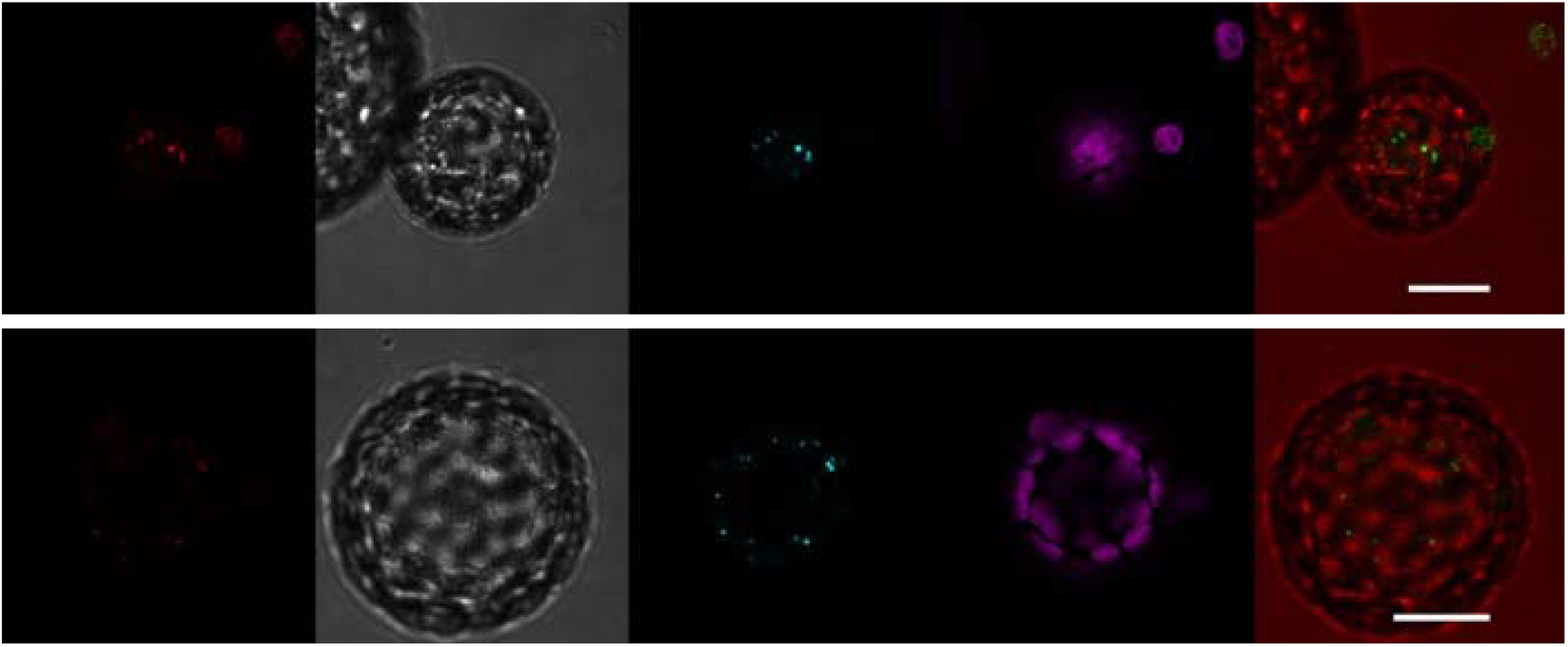
Recombinant catalase is located in the peroxisome. Co-localization of the florescent peroxisomal dye BODIPY with *H*. pylori catalase N-terminally tagged with M-Cherry in *N. benthamiana* protoplasts as imaged using confocal microscopy. Shown are M-Cherry (A), BODIPY peroxisomal dye (B), merged (C), chlorophyll fluorescence (D), and light (E) microscope image.

**Supplemental Table 1.** Various plant physiological and biochemical metrics were identified in WT, *cat2*-KO, Hp143, Hp432, and Hp615 lines. Leaf traits, photosynthetic metrics, *A*/*C*_*i*_ analysis, PIB analysis, Catalase Activity in WT, *cat2*-KO, Hp143, Hp432, and Hp615 lines. Shown are the means of 3-7 biological replicates with ± SE. Significant difference between the *cat2*-KO and individual heterologous lines are indicated by one (*p* ≤ 0.05), two (*p* ≤ 0.01), or three (*p* ≤ 0.001) red asterisk as determined by pairwise t-test with Bonferroni correction.

| Parameter | Unit | WT | *cat2*-KO | Hp143 | Hp432 | Hp615 |
| --- | --- | --- | --- | --- | --- | --- |
| **Leaf Traits** |  |  |  |  |  |  |
| Leaf Number | unitless | 37.3 ± 3.7 | 23 ± 1.4 | 26.8 ± 1.0 | 33 ± 0.6... | 47 ± 3.5... |
| Water Content | % | 92.4 ± 0.16 | 92.0 ± 0.37 | 91.7 ± 0.19 | 92.2 ± 0.06 | 92.4 ± 0.04 |
| *RGR* | mm<sup>2</sup> mm<sup>-2</sup> day<sup>-1</sup> | 0.254 ± 0.009 | 0.209 ± 0.009 | 0.224 ± 0.009 | 0.252 ± 0.008... | 0.256 ± 0.010... |
| *LMA* | g m<sup>-2</sup> | 21.1 ± 0.65 | 17.6 ± 0.85 | 19.2 ± 0.16 | 22.2 ± 0.41... | 22.9 ± 0.47... |
| *SLA* | m<sup>2</sup> kg<sup>-1</sup> | 47.6 ± 1.5 | 57.2 ± 2.9 | 52.0 ± 0.4 | 45.1 ± 0.9... | 43.7 ± 0.9... |
| **Photosynthesis** |  |  |  |  |  |  |
| $F_v / F_m$ | unitless | 0.751 ± 0.002 | 0.753 ± 0.003 | 0.762 ± 0.001. | 0.764 ± 0.002. | 0.764 ± 0.007. |
| *A* | μmol m<sup>-2</sup> s<sup>-1</sup> | 8.94 ± 0.788 | 6.29 ± 510 | 6.40 ± 0.832 | 8.11 ± 0.458... | 9.26 ± 0.451... |
| *g<sub>sw</sub>* | mol m<sup>-2</sup> s<sup>-1</sup> | 0.237 ± 0.0132 | 0.221 ± 0.0192 | 0.180 ± 0.0171 | 0.217 ± 0.0188 | 0.217 ± 0.0106 |
| *g<sub>m</sub>* (literature) | μmol m<sup>-2</sup> s<sup>-1</sup> Pa | 2.23 | 2.01 | 2.01 | 2.23 | 2.23 |
| *C<sub>i</sub>* | Pa | 32.6 ± 0.350 | 34.3 ± 0.252 | 34.3 ± 0.973 | 32.5 ± 0.603a. | 31.8 ± 0.379... |
| *R<sub>L</sub>* (literature) | μmol m<sup>-2</sup> s<sup>-1</sup> | 0.51 | 0.53 | 0.53 | 0.51 | 0.51 |
| *R<sub>D</sub>* | μmol m<sup>-2</sup> s<sup>-1</sup> | 1.36 ± 0.373 | 1.20 ± 0.324 | 1.59 ± 0.684 | 0.867 ± 0.364 | 0.626 ± 0.355 |
| $\Gamma^*$ (literature) | Pa | 4.482 | 5.827 | 5.827 | 4.482 | 4.482 |
| *v<sub>o</sub>* | μmol m<sup>-2</sup> s<sup>-1</sup> | 3.56 ± 0.394 | 2.47 ± 0.231 | 2.57 ± 0.376 | 3.19 ± 0.218. | 3.81 ± 0.246... |
| *v<sub>o</sub>* / *v<sub>c</sub>* | unitless | 0.314 ± 0.007 | 0.293 ± 0.004 | 0.295 ± 0.011 | 0.311 ± 0.007. | 0.325 ± 0.006... |
| **$A/C_i$ analysis** |  |  |  |  |  |  |
| *V<sub>c,max</sub>* | μmol m<sup>-2</sup> s<sup>-1</sup> | 37.8 ± 1.70 | 31.4 ± 6.19 | 34.8 ± 5.04 | 44.0 ± 5.79 | 46.3 ± 3.02 |
| *J* | μmol m<sup>-2</sup> s<sup>-1</sup> | 89.2 ± 4.25 | 79.8 ± 12.3 | 74.9 ± 9.1 | 99.6 ± 14.2 | 110.0 ± 7.6 |
| **PIB analysis** |  |  |  |  |  |  |
| *CO<sub>2</sub> Burst* | μmol m<sup>-2</sup> | 191 ± 22 | 404 ± 35.4 | 322 ± 43.2 | 196 ± 23.2... | 186 ± 12.5... |
| **Enzyme Activity** |  |  |  |  |  |  |
| Catalase 25°C | μmol H<sub>2</sub>O<sub>2</sub> m<sup>-2</sup> s<sup>-1</sup> | 457.7 ± 75.9 | 27.5 ± 7.81 | 44.7 ± 14.9 | 50.4 ± 11.0 | 78.7 ± 14.0. |
| **Safety Factor** |  |  |  |  |  |  |
| Catalase 25°C | unitless | 128.6 ± 21.3 | 11.1 ± 3.16 | 17.4 ± 5.81 | 15.8 ± 3.44 | 20.7 ± 3.67 |

**Supplemental Table 2.**
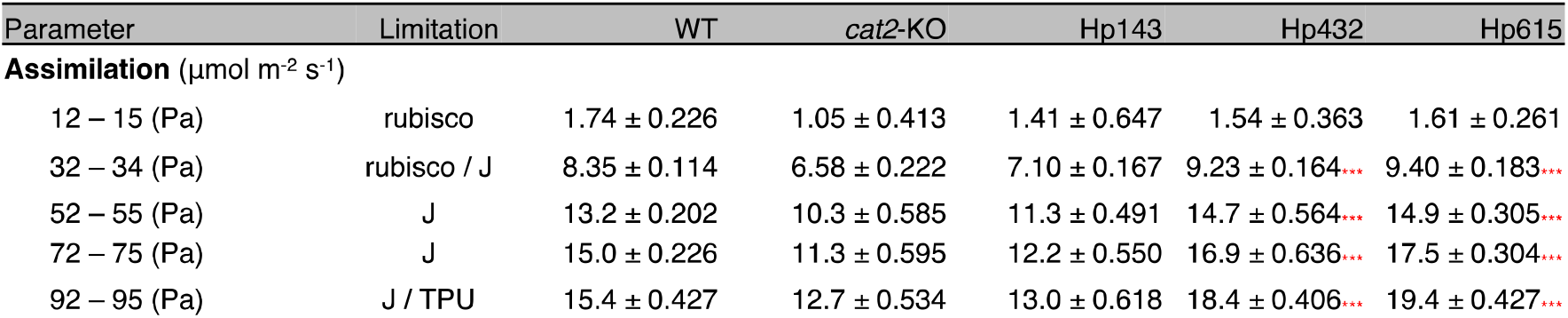
Net carbon fixation at various intercellular CO_2_ concentrations on the Dynamic Assimilation Technique (DAT) curve. Rates of net carbon fixation (*A*) were identified at 5 different intercellular CO_2_ concentrations in WT, *cat2*-KO, Hp143, Hp432, and Hp615 lines. Shown are the means of 3 biological replicates with ± SE. Significant difference between the *cat2*-KO and individual heterologous lines are indicated by one (*p* ≤ 0.05), two (*p* ≤ 0.01), or three (*p* ≤ 0.001) red asterisk as determined by pairwise t-test with Bonferroni correction.

